# SlowMoMan: A web app for discovery of important features along user-drawn trajectories in 2D embeddings

**DOI:** 10.1101/2022.08.23.505019

**Authors:** Kiran Deol, Griffin M. Weber, Yun William Yu

## Abstract

Nonlinear low-dimensional embeddings allow humans to visualize high-dimensional data, as is often seen in bioinformatics, where data sets may have tens of thousands of dimensions. However, relating the axes of a nonlinear embedding to the original dimensions is a nontrivial problem. In particular, humans may identify patterns or interesting subsections in the embedding, but cannot easily identify what those patterns correspond to in the original data. Thus, we present SlowMoMan (SLOW Motions on MANifolds), a web application which allows the user to draw a 1-dimensional path onto a 2-dimensional embedding. Then, by back-projecting the manifold to the original, high-dimensional space, we sort the original features such that those most discriminative along the manifold are ranked highly. We show a number of pertinent use cases for our tool, including trajectory inference, spatial transcriptomics, and automatic cell classification.

**Software availability:** https://yunwilliamyu.github.io/SlowMoMan/

**Code availability:** https://github.com/yunwilliamyu/SlowMoMan

## Introduction

The surge of big data in bioinformatics often means data sets may contain hundreds of thousands of dimensions/features [11, 14, 23]. Such data is impossible to visualize without low-dimensional embeddings. A number of linear and non-linear embedding algorithms exist, such as PCA [19], UMAP [3], and t-SNE [26]. Visual analysis tools can help users discover insights in their embeddings through visual enrichment of the embedding (such as by adding color maps, labels, annotations, etc.) [17]. Such tools allow users to manually identify areas of significance that may not be flagged as important by quantitative metrics alone. However, understanding the behaviour of the original features within such areas is nontrivial when the number of dimensions is too large to be manually examined. In particular, a user may wish to know if any of the original features exhibit special or notable behaviour within the area of interest.

Automatically determining discriminative features on labelled data (and then using those features for downstream tasks, such as prediction of labels on unlabelled data points) is a classic problem in machine learning, and has been solved in many ways using tools such as regression analysis [9], support vector machines (SVMs) [16], and deep neural networks [13]. However, in this paper we seek to identify those features which best determine/account for a particular spatial pattern in an embedding (as opposed to features which are most important in say, predicting the class of a point). For linear embedding methods like PCA, determining the most discriminative features is straightforward. For nonlinear methods, it is not clear how to determine which of the original features is most discriminative. As a proxy for discriminative ability over a particular spatial pattern, we propose a novel pipeline for ranking features by their periodicity along a user-defined path (1-dimensional manifold). In other words, we assume that features which best account for certain spatial patterns in an embedding must have some level of repeating structure along said manifold. Such a heuristic allows us to abstract away from the exact embedding algorithm used and also allow users to define completely custom paths. Additionally, our approach avoids making any decisions about which behaviours should be considered ‘discriminative’. Rather, we search for any kind of repeating structure.

To formalize the study of custom paths, we exploit the perspective of spatial analysis, i.e. studying the behaviour of features over a set of spatial parameters [25]. To illustrate, consider a two-dimensional city map with the two variables of rainfall and temperature overlaid. Analogously, we use a 2D embedding of high-dimensional data, and then seek to understand behavior of the features in the original space as a function of a some trajectory in the embedding space. The use of one-dimensional trajectories in our analysis enables us to leverage standard fast and efficient methods used in time series analysis to determine the importance of features.

Hence, in this manuscript, we seek to automatically extract features on non-categorical labels corresponding to user-specified trajectories/pseudo-time in datasets. What constitutes a notable area is dependent on the user’s domain knowledge, expectations, and purpose. For example, single cell trajectory inference seeks to identify continuous processes in single cell data, often corresponding to a notion of pseudotime in cell differentiation [12]. Often, such trajectories are near-to-obvious to the human eye when embedded in 2D, but the relationship between the 2D embedding and the original feature space may be complicated. Users can quickly identify potential trajectories in the embedded, but cannot easily link those trajectories back to feature space. Furthermore, users need to remain cognizant of spurious path-like structures in an embedding, which may appear due to the optimization criteria of various embeddings—t-SNE is well-known in certain parameter ranges to promote long thread-like regions, even when the underlying data do not strongly support it [27].

Thus, we present SlowMoMan, an in-browser tool which allows users to draw a custom, 1-dimensional path (a.k.a. manifold) over notable areas of their embedded data. The manifold therefore is an ordered list of points from the embedding. Using such a simple object means that our methodology does not rely on any assumptions about the choice of embedding method. By examining the visualization of the embedding and the distribution of the original features along possible trajectories in the embedding, users can quickly draw potential trajectories which they can compare and analyze. Once the user has drawn a path over a notable area, we run a nearest-neighbours algorithm on the points of the path to locate the nearest points in the embedding and then back-project those points to their original feature space (see. Fig. 1d). Along this back-projection, we examine the behaviour of each of the original features and for each of these features, we compute a user-chosen metric for discriminative ability.

**Figure 1.**
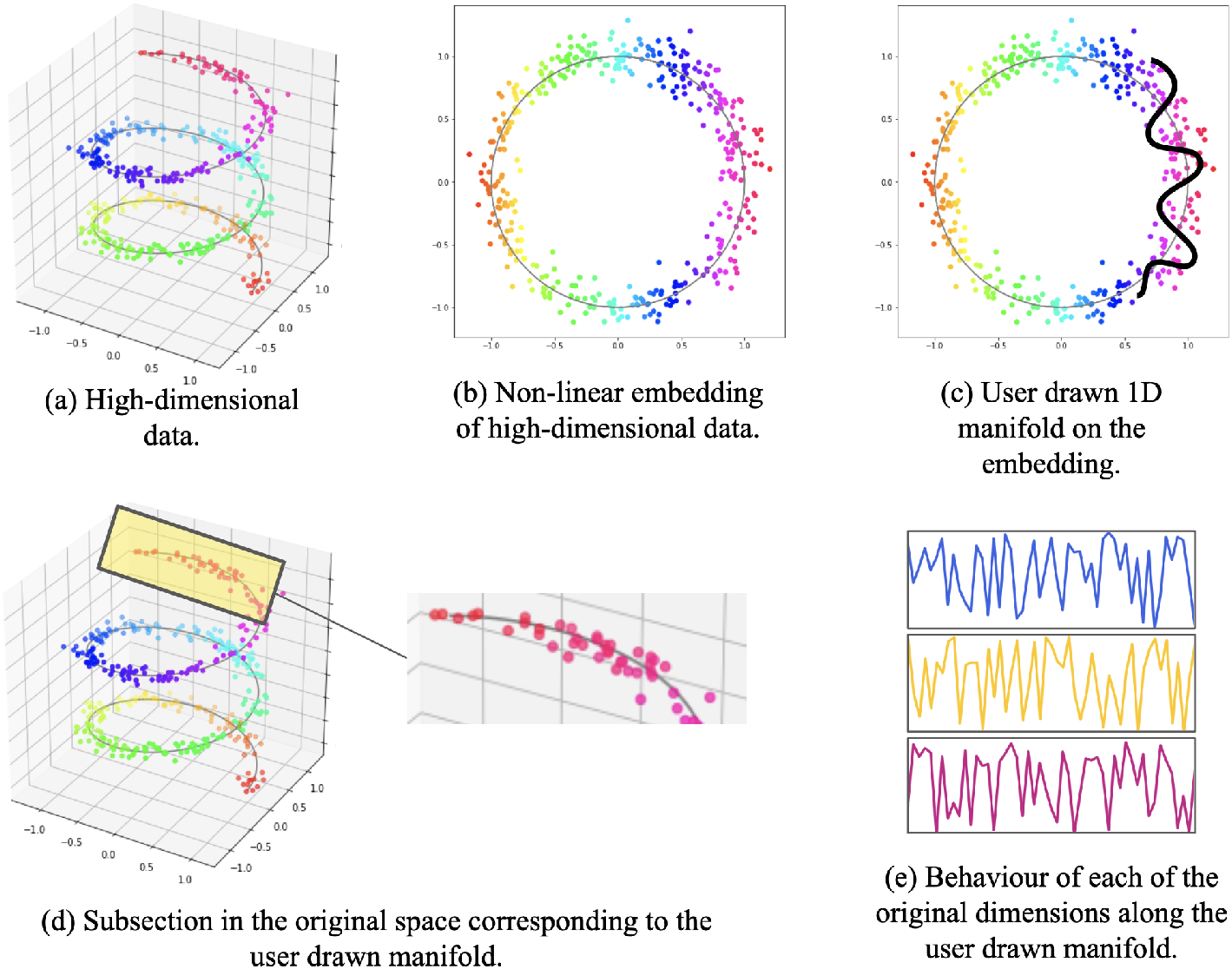
In most use cases, data with hundreds or thousands of dimensions would be best suited for SlowMoMan. (a) However, to visualize an example of high-dimensional data, we present a toy 3-dimensional dataset (the typical Swiss Roll dataset with 3 dimensions/features). (b) A non-linear embedding allows visualization in 2 dimensions, but it is not clear how these axes relate to the original features. (c) Users may use SlowMoMan to draw a path in the 2D embedding, which ideally corresponds to something close to a 1D manifold in the original space. (d) We can then cross-reference the points from the 2D embedding to their original representations in the high-dimensional space. (e) We analyze the behavior of the original features along that path, highlighting and sorting by the features that vary the slowest or are most autocorrelated.

Importantly, because the user has drawn a path, rather than provided a binary label, ‘discrimination’ here corresponds to measuring how important that feature seems to be to determining the position along the path (as opposed to how important a feature is in determining the label of some sample). Users may choose from autocorrelation or the harmonic sum of the magnitudes of the fast Fourier transform, two metrics we have found to be both quick-to-compute and that correspond well with a human intuition of path-importance. Based on the metric scores, we sort the original features. Using metrics to sort the original features provides a fast filter that does not impose any assumptions on the data set (as opposed to wrapper or embedded feature selection methods, which rely on generally expensive learning algorithms to rank features’ importance) [23]. The features ranked highly are then listed and their exact distributions along the manifold are displayed for the user, who can then manually validate the feature selection.

## Methodology

### Overview

SlowMoMan runs locally in-browser using the Javascript engine and relies on the user’s machine for all computation and storage. Thus, users do not need to install any additional software and can process their data sets without any third-party servers ever seeing their data. Once uploaded, the user will see the visualization of the embedded data. On this visual, the user can draw their own 1-dimensional manifold over any areas of interest (Fig. 1c). A nearest neighbours algorithm is then used to identify the points of the embedding which are nearest to the user’s custom manifold. Those points are then back-projected to their original space (Fig. 1d). Next, we observe the behaviour of each of the original dimensions along the back-projection of the user’s manifold (Fig. 1e).

Using the behaviour of each of the original dimensions along the back-projection of the user’s manifold, SlowMoMan will apply a number of possible metrics to sort the original dimensions, such that the dimensions most discriminative along the user’s manifold are ranked highest for further examination. Users will see a graph of the value of each of those recommended discriminative variables plotted against the time parameterization of the drawn trajectory. Users can also select specific variables to plot, if they have expert knowledge on which of the highest ranking variables are likely to be important.

### Canvas and back-projection

In our implementation, we used the D3 Javascript library to create a 700×700 SVG. The data is linearly rescaled to fit the 0 to 700 range. The choice of an SVG display enables more interactivity than a standard HTML canvas, including zooming and panning. However, the additional interactivity comes at a computational cost in that our display can handle roughly 7000 points before a notable lag is visible. We also include on the website an older non-vectorized version of SlowMoMan^1^ (v0) online that can handle ¿100,000 data points with 800 features or 3000 data points with 27,000 features, but that version lacks the quality-of-life features the reviewers requested.

Along the user-drawn path, we internally convert the path into 512 evenly spaced points on the canvas approximating that path. This is necessary because the mouse-over events vary across browsers, OS mouse sensitivity, and the mice themselves. If a user slowly drags a path, there will be many more mouse-over events than if a user draws the same path quickly; hence the normalization. The choice of *n* = 512 was largely arbitrary and users can modify this parameter via the “bin size” slider. Notably, for the FFT metric we restrict users to powers of 2 in order to take advantage of a more efficient FFT algorithm.

The choice of path can have significant implications for the final results. Crucially, our decision to use paths instead of clusters means that the order of points in the path plays an important role in the final computation. For example, a user can introduce artificial periodicity by drawing a line and then re-tracing that same line. Of course, this cannot modify the final rankings, as every feature would have the same level of artificial periodicity introduced. Given the inexact nature of path-drawing, SlowMoMan outlines the points used in the final computation in black, so users can see exactly which points have been selected and modify their path as needed.

For each of the 512 points along the path, we use a Quadtree (implemented with the D3 library) to find the nearest neighbour in the actual embedding. The Quadtree does not gurantee the true nearest neighbour is found, but its speed makes it a good choice for our application (as we may be doing this search 512+ times, depending on the bin size). Next, from the 512 embedding points, we simply grab their original representations in the high-dimensional space (back-projection). The actual feature ranking is computed on these high-dimensional points.

### Feature-ranking metrics

Depending on the data or the goals of the user, which features are most discriminative may vary. We explored a variety of possible metrics for users to choose from, though most performed fairly poorly (e.g. variance)—while variance is an accessible metric, its inability to account for the spatial/time-dependency along a path yields inconsistent results. The metrics that were most helpful were those that find periodicity along the path. As such, the metrics available in SlowMoMan include autocorrelation and the harmonic sum of the fast Fourier transform magnitudes (*FFT Score*).

#### Implementation details

All of SlowMoMan is implemented in client-side HTML5, Javascript, and CSS. The choice to use web technologies was made to take advantage of ready-to-use user interface elements, including the D3 visualization library and its pre-implemented zooming, panning, and dragging features. For the FFT computation, we use a 2014 version of the Project Nayuki Javascript FFT library [15], adapted by Cannam [4]. The autocorrelation computation does not use any libraries and is done directly in-app.

#### Autocorrelation

The autocorrelation of a variable x can intuitively be thought of as the level of correlation that x has with a delayed version of itself [10]. Within the domain of spatial analytics, the spatial autocorrelation is used to formalize the notion that areas near each other are more likely to be correlated. Thus, SlowMoMan uses the autocorrelation metric to test whether proximal points in the embedding are correlated within the original feature space. In other words, SlowMoMan uses autocorrelation to pick which features repeat themselves along the user-drawn path. Such periodicity may suggest that the feature is one of the determinants of the path. Formally, the autocorrelation is defined as

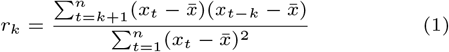

where 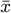 is the mean of x and k is the lag value. Thus, SlowMoMan computes *r*_*k*_ for each of the original dimensions along the back-projection of the embedding, using a default lag value of *k* = 22. The lag parameter controls the sampling rate of the feature. Thus at lag = 22, we compute the correlation between values 22 time steps apart. Since our default bin size is 512, we set the default lag to be the square root of 512. This of course is just a guess and the user can control the lag parameter via a slider.

However, one disadvantage of using autocorrelation is that we need to specify a lag value in advance. The choice of lag value *k* = 22 was found empirically to work reasonably well in our experimentation (close to the square root of the 512 path points), so that is the default lag, but that varies by application. Due to the need to set a parameter, autocorrelation should only be used by advanced users with a reasonable guess as to the lag that should be set.

#### Fast Fourier Transform

That of course leads us to desire a parameter-less way to find features that have some periodicity. Thanks to the decision to rely on a single parameter in our spatial analysis, we can leverage well-known tools used in time series analysis. To this end, we turn to standard signal processing tools and the Fourier transform. The fast Fourier transform is an efficient algorithm for computing the discrete Fourier transform [6]. The discrete Fourier transform decomposes a sequence of values into a sum of discrete sinusoidal waves, each with a distinct magnitude and frequency. Formally, if we have a sequence of k values *{x*_*n*_*}*, we can approximate it with another sequence *{F*_*m*_*}* of sinusoids.

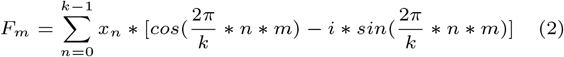

Each of these new sinusoids will have a respective amplitude, which is the absolute height of the sinusoid. Across all of the amplitudes *m*_*n*_ of each of the sinusoids for a specific sequence *{x*_*n*_*}*, we can take a harmonic sum S over descending *m*_*n*_

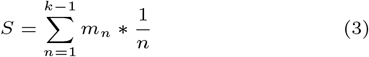

which is then used as the custom “FFT Score” metric. Notice the metric is maximized when a feature can be decomposed into sinusoids with large amplitudes - as opposed to features with many small amplitude sinusoids, which may be indicative of noise. In other words, the harmonic sum inherently lends greater weight to the sinusoids with lower frequencies— although, we naturally omit the constant *m*_0_ term as we are only interested in periodic behavior. Intuitively, we are not as interested in high-frequency variables because they are not strongly correlated with position on our path. Instead, low-frequency variables (i.e. slow motions) are features where if you know their values, you can roughly predict your position along the path. This allows us to simultaneously capture periodicity at all scales, while more heavily weighting longer periods, overcoming the shortcoming of having to set a lag parameter in autocorrelation.

## Evaluation

We demonstrate the utility of SlowMoMan through case studies and empirical analysis. For all use cases presented below, we have made available associated data and trajectory information on the SlowMoMan website (see “Use Cases” and from there “Google Drive artifacts”). In the first task, our prototypical use case, we show how SlowMoMan can be used to help interpret trajectories drawn on a 2D embedding. In this task, we consider two trajectories, where the features are actual genes in single-cell transcriptomic data. In the second task, we demonstrate using SlowMoMan on a more classic task related not to trajectories, but to simple clusters instead—we find important features disambiguating between two clusters of spatial transcriptomic cell data. In the last task, we use SlowMoMan to better characterize the nature of errors in automatic cell classification using single-cell transcriptomic data.

### Trajectory Inference/Pseudotemporal Ordering

We apply SlowMoMan to datasets taken from Eulenberg et al. [8] (Fig. 2) and Paul et al. [18] (Fig. 3). Physical limitations in the technologies used to obtain gene expression data often make it impossible to track dynamic processes in single cells, such as the cell cycle or cell differentiation. Trajectory inference methods seek to infer such processes by analyzing similarities in the gene expression of cells to order them along a continuous interval. A large number of algorithms exist for such a purpose and Saelens et al. [20] benchmark 45 of these methods on their accuracy, scalabilty, stability, and usability. SlowMoMan inherently complements such methods due to its focus on trajectories, as opposed to standard tools which often focus on analyzing entire clusters. In particular, SlowMoMan may be used to identify important features along a given trajectory or to quickly explore alternative, user-defined trajectories.

**Figure 2.**
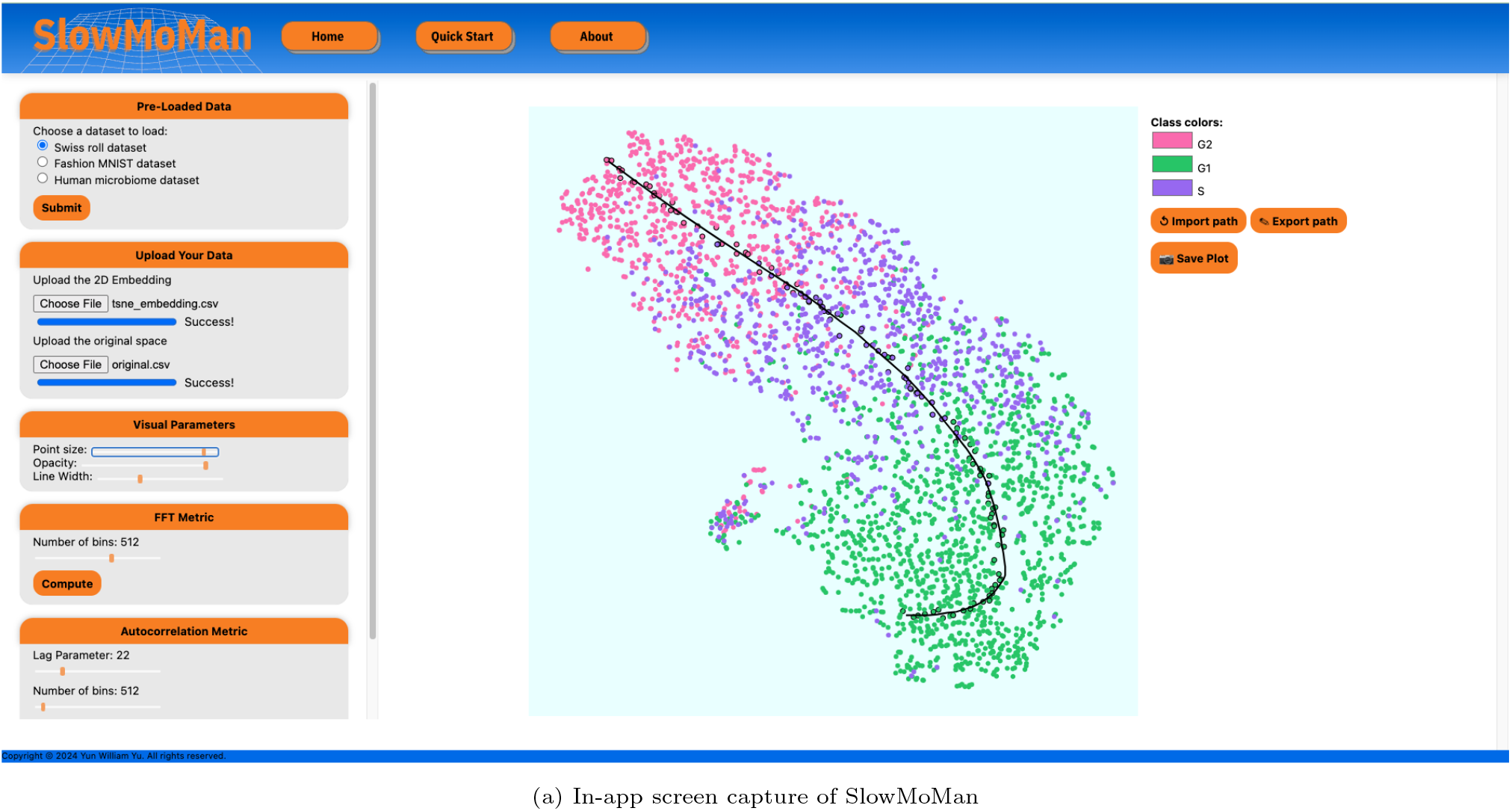
SlowMoMan identifies the most informative nodes within a neural network trained to process images of Jurkat cells in different stages of the cell cycle [8]. The t-SNE embedding of the neural network activation space representations of each cell is presented near the center of the screen capture. Notice the clusters already have a fairly clear trajectory and are indeed in their correct order. The top 7 most important nodes identified by SlowMoMan are displayed in the left half of the lower two boxes. The behaviour of the top 7 nodes along the path of the trajectory is displayed in the right half of the lower two boxes. Notably, all of the nodes move in a fairly similar pattern, as to be expected with a good trajectory.

**Figure 3.**
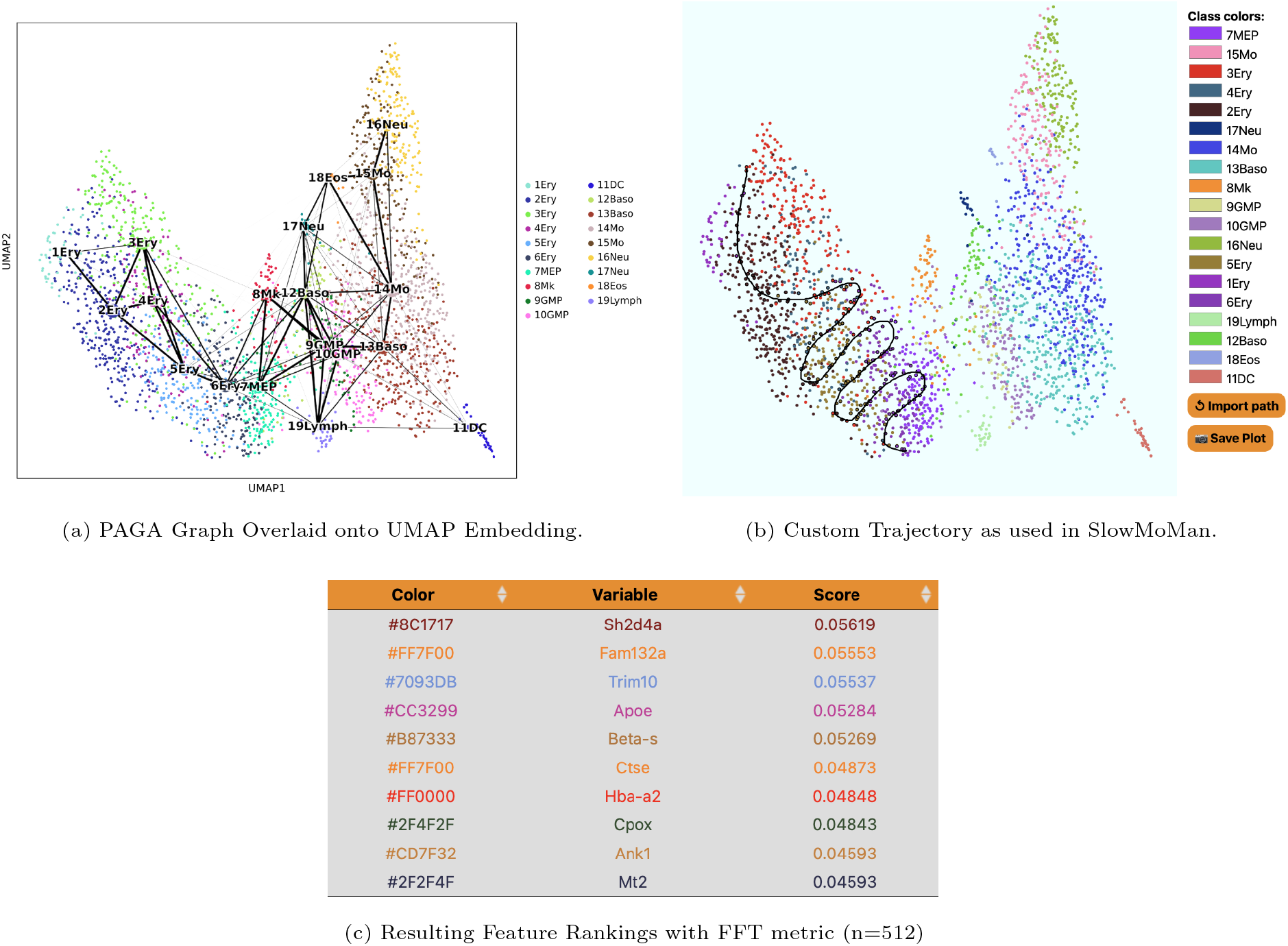
The results of the PAGA algorithm on the hematopoiesis data of Paul et al, 2015 [18] highlight strong connectivities between clusters 1-6. We use SlowMoMan to explore important features along a custom manifold between clusters 1-7. (a) The UMAP embedding of the hematopoiesis data of Paul et al, 2015 [18] with the PAGA connectivities between clusters represented as black edges. Thicker edges indicate stronger connectivities. Clusters correspond to cell type, with labels corresponding to those of Paul et al, 2015 [18]. Note also the prominent connectivities between clusters 1-7. (b) A screen capture of SlowMoMan with the user-drawn manifold used in the analysis.

#### Trajectory Inference in Hepatoblast Differentiation of Mouse Liver Cells

Data collected by Yang et al. [31] was retrieved from a Kaggle dataset [5]. The raw data contains 24,748 gene counts across 447 liver cells obtained from mice. Each cell is labelled by its embryonic date, starting from embryonic day 10.5 (E10.5) to embryonic day 17.5 (E17.5), with E16.5 excluded. For our analysis (Fig. 4), we changed the labelling of embryonic days to a simple scalar where E10.5 became 1, E11.5 became 2, and so fourth (although E16.5 is not present in the dataset, E17.5 is still labelled 8). Although the authors used PCA to obtain their 2D embedding, we chose to use UMAP. The linearity of PCA makes it a feature importance method in its own right [21] (thus making its usage with SlowMoMan somewhat redundant). Non-linear methods like UMAP, which do not have clear means of identifying the contribution of each feature, are better suited to SlowMoMan. As a simple sanity check, we demonstrate a run of SlowMoMan with the FFT metric.

**Figure 4.**
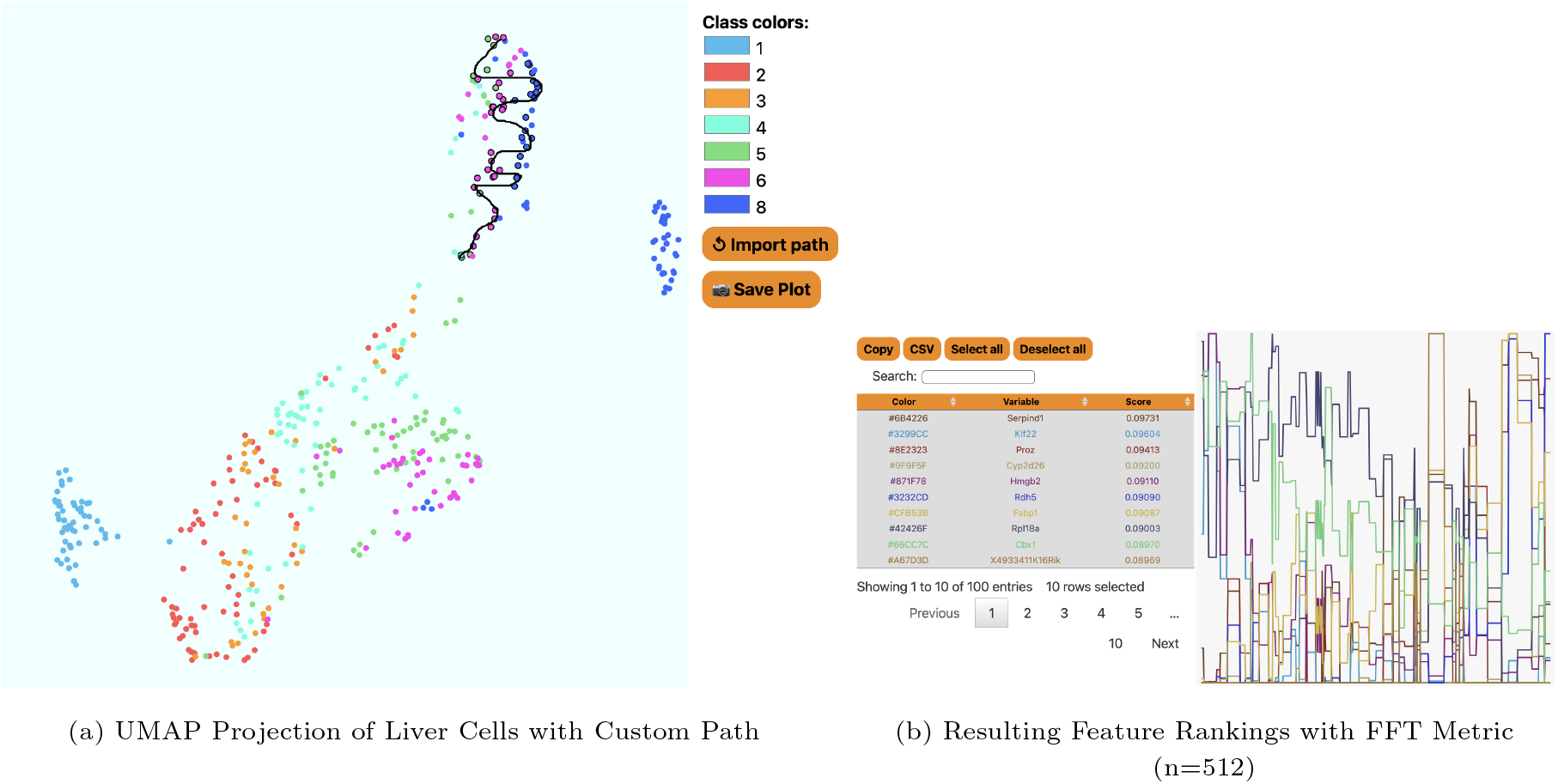
In (a) we present the 2D embedding displayed by SlowMoMan for mouse liver cells, where the labels on the right correspond to embryonic date. Notice the trajectory is drawn over a fairly isolated cluster of cells containing only days 4, 5, 6, and 8. In (b) we show the final feature rankings as given by SlowMoMan. Note that only the top 10 features are actually displayed.

For the first run, we present an irregular scrawl drawn over an interesting cluster of the UMAP embedding (see Fig.4b). This cluster is off to the top right of the “main” cluster seen below it. This cluster is particularly interesting in that it only contains days 5, 6, and 8 (recall day 7 is not in the dataset) and each of those days present themselves in roughly vertical lines in roughly the correct order (days 4 and 5 are intermixing, but days 6 and 8 are quite distinct). The choice to use a “squiggle” shape comes from the fact that since each day presents roughly vertically, the overall progression from day 4 to 5, and 5 to 6, etc. is therefore horizontal. However, since this cluster is not very wide, a single horizontal line would be not be representative of the entire cluster. Thus, the squiggle manages to create multiple horizontal lines on the cluster. After drawing this, we compute the FFT metric. Recall that there are nearly 25,000 genes and we rely on the FFT metric to return a list of the top 100 genes who present periodic behaviour along our manifold. The analysis of the genes marked significant by the FFT metric is of course limited to the amount of domain knowledge and time for exploration that one has available.

We now discuss a few notable genes flagged by the FFT metric. Ranked 1st was SERPIND1, which is known to have highly biased expression in the liver [22]. Ranked 4th was CYP2D26 which the original authors identified as notable among the E15.5 and E17.5 hepatocytes (see fig 2 of [31]). This aligns well with the fact that the cluster being analyzed here contained days 6 and 8 most prominently. Ranked 7th was FABP1 (fatty acid binding protein 1), which is known to have expression in the liver as it encodes the for fatty acid binding protein of the liver [22]. Although this evaluation is inherently limited by our domain knowledge and exploration, the ability to flag such genes as significant (out of 25,000) is remarkable.

#### Trajectory Inference in Hematopoiesis of Mouse Bone Marrow Cells

Data collected by Paul et al. [18] was retrieved through the *paul15()* method of [28]. The raw data contains 3,451 gene counts across 2,730 labelled bone marrow cells of mice. The clusters correspond to cell type. Numbers are prefixed to the cluster labels to distinguish between cells that may belong to the same cell type, but are at different levels of hematopoietic differentiation. In order to identify potential trajectories we use PAGA [29], one of the best overall trajectory inference methods in [20]. From a k-nearest neighbours (kNN) graph of single-cells and the associated clusters (partitions) of these cells, PAGA will compute a new graph representing the connectivities of the partitions. Obtaining the data, preprocessing, and running PAGA on this dataset were all accomplished through the Python package Scanpy, which contains methods for trajectory inference [28]. In particular, we use UMAP with the euclidean distance metric to compute a single-cell connectivity graph with a neighbourhood size of 20. We use the existing clusters of Paul et al. [18] as the partitions. We then run PAGA to obtain the graph of connectivities between partitions, as shown in Fig. 3.

A notable set of connectivities are those among clusters 1 to 7, which Paul et al. [18] hypothesize represent a continuum of erythrocyte differentiation. SlowMoMan is used to draw a trajectory between these clusters (see Fig.3) and using the FFT score to sort, obtain a list of the 100 most important features out of 3451. Notably, a number of key transcription factors identified in Fig. 1C of Paul et al. [18] are ranked within the top 20 genes of this trajectory by SlowMoMan, includinig *Apoe, CPOX, Hba-a2*, and *Car2*. Although not previously associated with erythropoiesis, Paul et al. [18] noted the significance of *Cited4* in clusters 1-7. Interestingly, SlowMoMan ranks *Cited4* at 76th. Although SlowMoMan was able to identify a number of key genes, due to these genes being spread across the list of top 100 features, a user without prior knowledge may not realize their significance. Indeed, users with prior knowledge as which features are most important may focus on using SlowMoMan to experiment with possible trajectories that capture as many of the important features as possible, as illustrated in the current example. In contrast, users without prior expectations on the features may instead use SlowMoMan is to gather a list of important features along a specific trajectory which may be investigated further with more conclusive methods.

### Spatial Transcriptomics

We apply SlowMoMan to a Human Lymph Node Spatial Gene Expression Dataset by Space Ranger 1.0.0, as provided by 10x Genomics [1] (Fig. 5). A recent advancement in bioinformatics is the ability to associate gene expression data with its spatial distribution in a sample. Specifically, spatial transcriptomics enables researchers to locate the position of individuals cells (and the respective gene expression data of each cell) in a sample. Spatial transcriptomics is a strong candidate field for SlowMoMan, as clusters in a tissue sample are unlikely to be perfectly distinct. Indeed, while the clusters of the corresponding embedding of the cells of tissue sample cells tend to remain fairly coherent, the clusters in the tissue sample itself are often intermingled are broken into a number of subclusters, as illustrated in 5. In such cases, analyzing the features of entire clusters would not be as informative as analyzing the individual subsections of clusters and comparing them.

**Figure 5.**
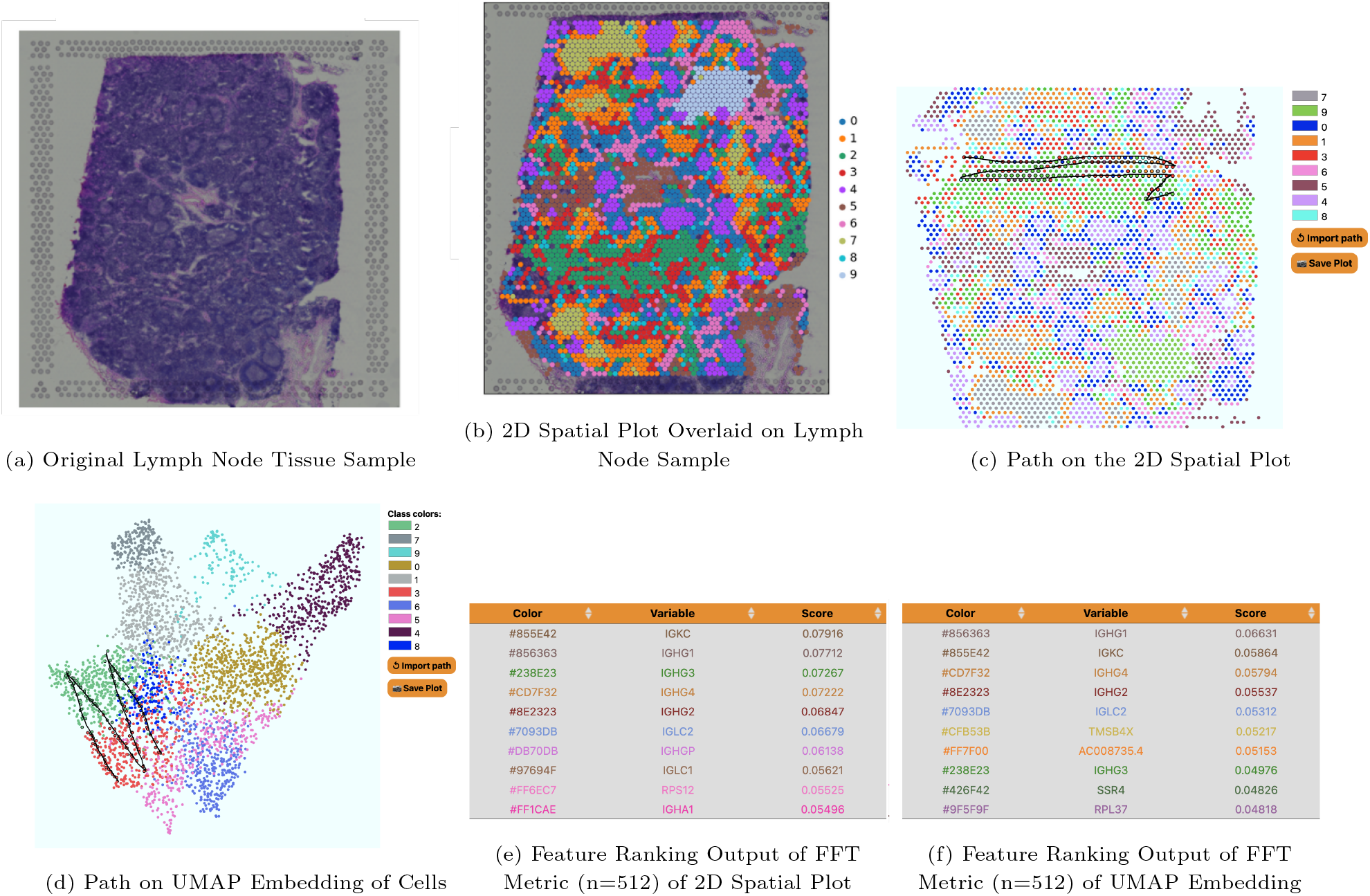
SlowMoMan applied to a spatial transcriptomics dataset to determine important features distinguishing between two nearby clusters. (a) The original image of the human lymph node tissue sample [1]. (b) The human lymph node tissue sample with the cells and their respective UMAP classes overlaid. (c) The path drawn on the 2D spatial plot as used SlowMoMan. (d) The resulting feature ranking based on the path drawn in (c). (e) The path drawn on the 2D UMAP embedding as used SlowMoMan. (f) The resulting feature ranking based on the path drawn in (e). Notice that many of these features were also ranked highly in the FFT results on the 2D spatial plot.

Thus, we apply SlowMoMan to a Human Lymph Node Spatial Gene Expression Dataset, provided by 10x Genomics [1] (Fig. 5). In particular, 10x Genomics used their Visium Spatial Gene Expression tool to analyze a sample of human lymph node tissue. Obtaining the data, preprocessing, visualization, and running the embedding algorithm were all accomplished through the Python package Scanpy, which also contains methods for spatial transcriptomics [28].

This dataset from 10x Genomics contains the spatial coordinates of each of the individually identified cells in the tissue sample, along with the gene expression levels of each of these cells. From the gene expression data, we use UMAP on a 50-dimensional PCA embedding to compute a graph with neighbourhoods of size 15. This neighbourhood graph is then used in the Leiden clustering algorithm of [24] to identify 10 within the data. Although these clusters are distinct (see 5), we cannot yet make any claims as to what known lymph cell subtypes these clusters may translate to. However, we can identify patterns between the spatial distribution of the cells and their UMAP distribution and use SlowMoMan to explain the genetic underpinnings for such patterns.

One notable observation is that clusters 2 and 3 (green and red) not only exist beside each other in the UMAP embedding, but are also frequently seen together in the tissue sample, with red appearing almost as a border to the green sections. In contrast to the embedding, the red cluster is broken up into many separate, smaller subclusters throughout the tissue sample. Computing the feature importance of the red cluster in its entirety would fail to capture any behaviour unique to aforementioned area of the red cluster which borders the green cluster. Thus, we first use SlowMoMan without considering any subclusters and analyze the red and green clusters in the UMAP embedding. Then, we compare the results to SlowMoMan being run on the aforementioned subcluster of the spatial distribution. Notice in Fig. 5 that the path drawn goes back-and-forth between the two clusters, unlike in the previous example; this artifically creates the periodicity that SlowMoMan tries to detect, allowing it to find not just discriminating features for paths, but also clusters.

Between the two runs of SlowMoMan shown in Fig. 5, we used the FFT score to obtain the top ten most important genes in both instances and found that genes IGHG1, IGHG2, IGHG3, IGHG4, IGLC2, and IGKC were all identified among the top 10 most important genes (out of 19,686) in both the trajectory drawn on the UMAP embedding and the trajectory drawn on the spatial distribution. Notably, the Entrez Gene summary for each of these genes predicts that they are important to immunoglobulin receptor binding. Immunoglobulins are produced by B-lymphocytes, which are located primarily in the cortex of the lymph nodes. Given that the red and green clusters were observed mainly in one region of the tissue sample, we could the explain the importance of IGHG1, IGHG3, IGHG4, and IGKC as the result of the cells of the red and green clusters being located in the cortex of the lymph node tissue sample.

### Automatic Cell Classification

We apply SlowMoMan to a single-cell transcriptomic dataset from [30] (Fig. 6). In single cell analysis, data sets are often too large for manual annotation and methods for automated classification of cell types are a promising alternative. A number of challenges arise when using such methods, however, such as choosing which method to use or evaluating the accuracy of a given method. SlowMoMan augments such methods by allowing users to interpret regions where multiple classification methods may differ, or by interpreting regions where a classification method fails.

**Figure 6.**
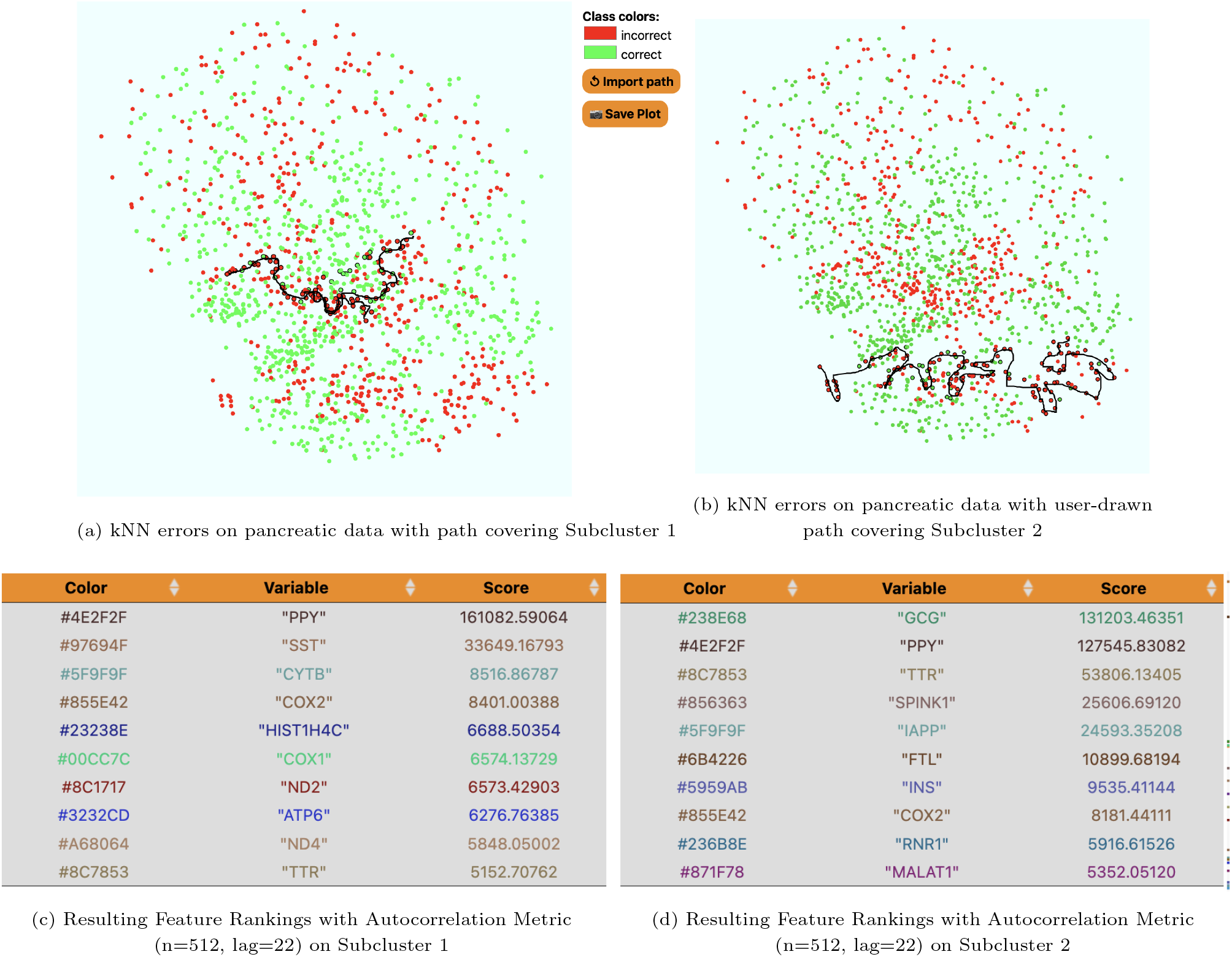
SlowMoMan is used to understand the types of errors made by the kNN algorithm (with a neighbour size of 9) on an automatic cell classification task in single-cell transcriptomics [30]. We use UMAP to obtain a 2-dimensional embedding. The clusters represent which cells were classified correctly vs. incorrectly by the trained kNN model. Of the 581 incorrectly labelled cells, 78% were beta cells, 14% were gamma cells, and 8% were delta cells. None of the incorrectly identified cells were alpha cells. (a) A screen capture of SlowMoMan with the user-drawn trajectory over a fairly distinct subcluster of errors. Notable genes identified along this trajectory were NPY, PPY, and ZFP36. (b) Similar to (a), but with a user-drawn trajectory covering a different, but also fairly distinct subcluster of errors. Notable genes identified along this trajectory were G6PC2 and SCGB2A1.

For example, the pancreatic data obtained by [30] examines the transcriptomes of 1492 islet cells across 18 donors and includes four classes of islet cells: alpha, beta, gamma, and delta. As part of a survey of automatic cell classification methods, [2] apply a number of classification methods to this dataset, including the kNN algorithm (k = 9). Replication of these methods found that kNN had an accuracy score of 59.10%, supporting the results of [2]. SlowMoMan augments this method by enabling the interpretation of regions where kNN fails.

One explanation for the poor performance of the kNN algorithm was the imbalance in data: 60% were alpha cells, 32% were beta cells, 5% were gamma cells, and 3% were delta cells. This would also explain why all of the incorrectly identified cells were either beta, gamma, or delta cells. However, we also use SlowMoMan to understand the possible genomic causes for these errors. In particular, we run SlowMoMan on two large and fairly distinct subclusters of kNN errors (see 6). Akin to the previous example, analyzing the feature importances across the entire cluster of errors would fail to capture any information unique to the two identified subclusters. Thus, we perform two runs of SlowMoMan, one for each of the previously identified subclusters of errors.

For both subclusters, we manually examined the final results and simply researched a few of the top 100 genes. Despite the inexactness of this approach, its exploratory and casual nature demonstrates our typical use-case.In the first subcluster (Fig. 6a), we identified 3 genes of interest: NPY, PPY, and ZFP36. In the second subsection, (Fig. 6b), we identified 2 genes of interest: G6PC2 and SCGB2A1. We note that in both subclusters, the aforementioned genes were all within the top

100 most important genes (out of 33890) of their respective trajectories. We also note that in this case, sorting with the autocorrelation metric led to more notable genes being identified than with the usual FFT score. The PPY gene is a known marker gene of gamma cells. Thus, its importance is to be expected, given the inability of the kNN model to recognize any of the gamma cells. The importance of the remaining four genes, however, can be explained by the discovery of four beta cell sub-types by [7]. Indeed, they identify the four subtypes as *β*_1_, *β*_2_, *β*_3_, *β*_4_. Within which there is the *ST* 8*SIA*1^+^ *β*_3_/*β*_4_ subtype, *ST* 8*SIA*1^−^ *β*_1_/*β*_2_ subtype, *CD*9^+^ *β*_2_/*β*_4_ subtype, and the *CD*9^−^ *β*_1_/*β*_3_ subtype. Notably, NPY and ZFP36 were found by [7] to be differentially expressed between the *CD*9^−^ *β*_1_/*β*_3_ and *CD*9^+^ *β*_2_/*β*_4_ subtypes. In particular, NPY had larger expression levels in the *CD*9^−^ *β*_1_/*β*_3_ subtype, whereas ZFP36 had larger expression levels in the *CD*9^+^ *β*_2_/*β*_4_ subtype. Thus, although the cells in the first subsection of errors (Fig. 6a), were primarily beta cells, the possibility of multiple subtypes of beta cells existing in that area may have contributed to the poor results obtained by the kNN model when classifying beta cells. Similarly, in the second subsection analyzed (Fig. 6b), the genes G6PC2 and SCGB2A1 were also identified by [7] as being differentially expressed between the *ST* 8*SIA*1^+^ *β*_3_/*β*_4_ and *ST* 8*SIA*1^−^ *β*_1_/*β*_2_ subtypes. Therefore, even with an imbalance in the data set, the analysis with SlowMoMan suggests the possibility that the unspecified subtypes within the beta cells may have contributed to the poor performance of the kNN model on classifying beta cells.

### Runtime Evaluation

We provide runtime profiling of SlowMoMan in Table 1 Profiling was done with 8 GB of RAM and the 2.3 GHz Dual-Core Intel Core i5 processor (2017). The time to load column represents the total time to load the embedding and original space. The time to compute metric represents the time taken to compute the respective metric of that use case.

**Table 1.**
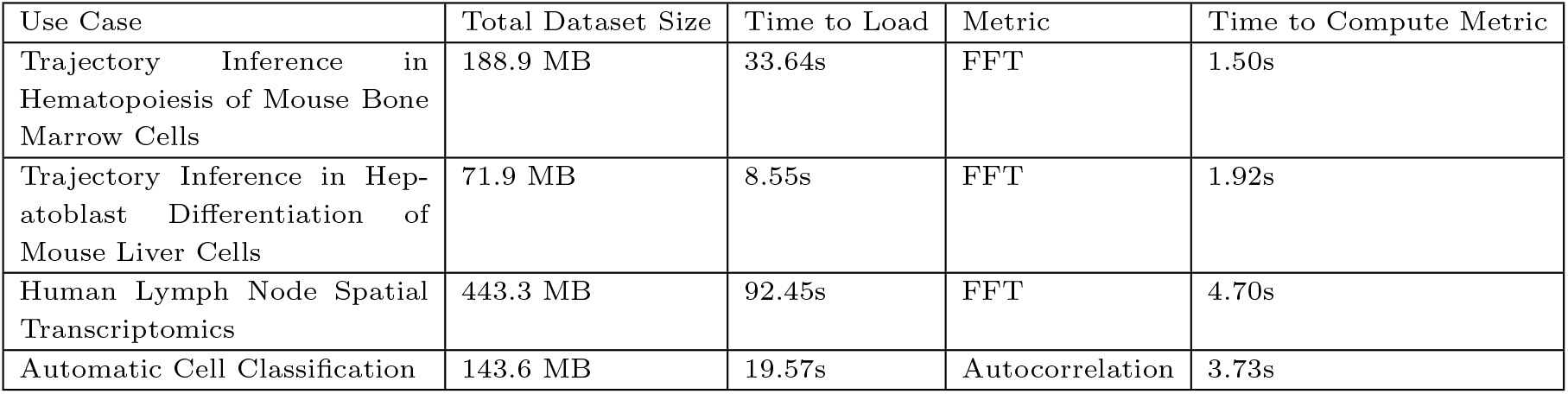
Runtime benchmarks.

## Conclusion

In this manuscript, we presented a new interactive web app, SlowMoMan, for visualization and exploration of important variables in low-dimensional embeddings. This web app runs entirely client-side using Javascript, HTML 5, and CSS, and allows for real-time usage on commodity laptops. By back-projecting user-drawn paths and ranking variables by proxies for their periodicity, users are able to identify the original features that are most informative for those paths.

The major advance of our work is in allowing users to draw paths, rather than simply apply categorical labels, which we showed in our two trajectory analysis examples. As shown in our latter two examples, users are still able to examine features important for binary classifications by simply drawing appropriate paths doubling back on themselves. However, the paths lend themselves particularly well in cases where there is some underlying notion of paths and time, such as in cell differentiation and cell cycles. We hope tools like SlowMoMan will assist scientists in interpreting and analyzing low-dimensional embeddings.

## Author contributions statement

Y.W.Y. and G.W. conceived the project. Y.W.Y. and K.D. wrote the software. K.D. conducted the experiments.

## Acknowledgments

K.D. was supported by the University of Toronto Data Sciences Institute as a Summer Undergraduate Data Science (SUDS) Research Scholar. We acknowledge the support of the Natural Sciences and Engineering Research Council of Canada (NSERC) (NSERC grant RGPIN-2022-03074) and DND/NSERC Discovery Grant Supplement DGDND-2022-03074.

## Footnotes

1 https://github.com/yunwilliamyu/SlowMoMan_v0

## References

1. 10x Genomics. Human lymph node spatial gene expression dataset by space ranger 1.0.0. https://support.10xgenomics.com/spatial-gene-expression/datasets/1.0.0/V1_Human_Lymph_Node? Accessed: 2022-08-01.

2. Tamim Abdelaal, Lieke Michielsen, Davy Cats, Dylan Hoogduin, Hailiang Mei, Marcel J. T. Reinders, and Ahmed Mahfouz. A comparison of automatic cell identification methods for single-cell rna sequencing data. Genome Biology, 20(1):194, Sep 2019.

3. Etienne Becht, Leland McInnes, John Healy, Charles-Antoine Dutertre, Immanuel WH Kwok, Lai Guan Ng, Florent Ginhoux, and Evan W Newell. Dimensionality reduction for visualizing single-cell data using umap. Nature biotechnology, 37(1):38–44, 2019.

4. Chris Cannam. Javascript dsp tests. https://code.soundsoftware.ac.uk/projects/js-dsp-test/repository/show/fft. Accessed: 2019-08-01.

5. Alexander Chervov. scrna-seq trajectory inference, 2022.

6. James W Cooley and John W Tukey. An algorithm for the machine calculation of complex fourier series. Mathematics of computation, 19(90):297–301, 1965.

7. Craig Dorrell, Jonathan Schug, Pamela S. Canaday, Holger A. Russ, Branden D. Tarlow, Maria T. Grompe, Tamara Horton, Matthias Hebrok, Philip R. Streeter, Klaus H. Kaestner, and Markus Grompe. Human islets contain four distinct subtypes of β cells. Nature Communications, 7(1):11756, Jul 2016.

8. Philipp Eulenberg, Niklas Köhler, Thomas Blasi, Andrew Filby, Anne E Carpenter, Paul Rees, Fabian J Theis, and F Alexander Wolf. Reconstructing cell cycle and disease progression using deep learning. Nature Communications, 8(1):463, September 2017.

9. Rudolf J Freund, William J Wilson, and Ping Sa. Regression analysis. Elsevier, 2006.

10. John A Gubner. Probability and random processes for electrical and computer engineers. Cambridge University Press, 2006.

11. Melanie Hilario and Alexandros Kalousis. Approaches to dimensionality reduction in proteomic biomarker studies. Briefings in bioinformatics, 9(2):102–118, 2008.

12. Zhicheng Ji and Hongkai Ji. Tscan: Pseudo-time reconstruction and evaluation in single-cell rna-seq analysis. Nucleic acids research, 44(13):e117–e117, 2016.

13. Yann LeCun, Yoshua Bengio, and Geoffrey Hinton. Deep learning. nature, 521(7553):436–444, 2015.

14. Jason H Moore, Folkert W Asselbergs, and Scott M Williams. Bioinformatics challenges for genome-wide association studies. Bioinformatics, 26(4):445–455, 2010.

15. Project Nayuki. Free small fft in multiple languages. https://www.nayuki.io/page/free-small-fft-in-multiple-languages. Accessed: 2022-08-01.

16. William S Noble. What is a support vector machine? Nature biotechnology, 24(12):1565–1567, 2006.

17. Luis Gustavo Nonato and Michäel Aupetit. Multidimensional projection for visual analytics: Linking techniques with distortions, tasks, and layout enrichment. IEEE Transactions on Visualization and Computer Graphics, 25(8):2650–2673, 2019.

18. Franziska Paul, Ya’ara Arkin, Amir Giladi, Diego Adhemar Jaitin, Ephraim Kenigsberg, Hadas Keren-Shaul, Deborah Winter, David Lara-Astiaso, Meital Gury, Assaf Weiner, Eyal David, Nadav Cohen, Felicia Kathrine Bratt Lauridsen, Simon Haas, Andreas Schlitzer, Alexander Mildner, Florent Ginhoux, Steffen Jung, Andreas Trumpp, Bo Torben Porse, Amos Tanay, and Ido Amit. Transcriptional heterogeneity and lineage commitment in myeloid progenitors. Cell, 163(7):1663–1677, 2015.

19. Markus Ringnér. What is principal component analysis? Nature biotechnology, 26(3):303–304, 2008.

20. Wouter Saelens, Robrecht Cannoodt, Helena Todorov, and Yvan Saeys. A comparison of single-cell trajectory inference methods. Nature Biotechnology, 37(5):547–554, May 2019.

21. Fengxi Song, Zhongwei Guo, and Dayong Mei. Feature selection using principal component analysis. In 2010 International Conference on System Science, Engineering Design and Manufacturing Informatization, volume 1, pages 27–30, 2010.

22. Gil Stelzer, Naomi Rosen, Inbar Plaschkes, Shahar Zimmerman, Michal Twik, Simon Fishilevich, Tsippi Iny Stein, Ron Nudel, Iris Lieder, Yaron Mazor, Sergey Kaplan, Dvir Dahary, David Warshawsky, Yaron Guan-Golan, Asher Kohn, Noa Rappaport, Marilyn Safran, and Doron Lancet. The genecards suite: From gene data mining to disease genome sequence analyses. Current Protocols in Bioinformatics, 54(1):1.30.1–1.30.33, 2016.

23. F William Townes, Stephanie C Hicks, Martin J Aryee, and Rafael A Irizarry. Feature selection and dimension reduction for single-cell rna-seq based on a multinomial model. Genome biology, 20(1):1–16, 2019.

24. V A Traag, L Waltman, and N J van Eck. From louvain to leiden: guaranteeing well-connected communities. Scientific Reports, 9(1):5233, March 2019.

25. D. J. Unwin and L. W. Hepple. The statistical analysis of spatial series. Journal of the Royal Statistical Society. Series D (The Statistician), 23(3/4):211–227, 1974.

26. Laurens Van der Maaten and Geoffrey Hinton. Visualizing data using t-sne. Journal of machine learning research, 9(11), 2008.

27. Martin Wattenberg, Fernanda Viégas, and Ian Johnson. How to use t-sne effectively. Distill, 1(10):e2, 2016.

28. F. Alexander Wolf, Philipp Angerer, and Fabian J. Theis. Scanpy: large-scale single-cell gene expression data analysis. Genome Biology, 19(1):15, Feb 2018.

29. F. Alexander Wolf, Fiona K. Hamey, Mireya Plass, Jordi Solana, Joakim S. Dahlin, Berthold Göttgens, Nikolaus Rajewsky, Lukas Simon, and Fabian J. Theis. Paga: graph abstraction reconciles clustering with trajectory inference through a topology preserving map of single cells. Genome Biology, 20(1):59, Mar 2019.

30. Yurong Xin, Jinrang Kim, Haruka Okamoto, Min Ni, Yi Wei, Christina Adler, Andrew J. Murphy, George D. Yancopoulos, Calvin Lin, and Jesper Gromada. Rna sequencing of single human islet cells reveals type 2 diabetes genes. Cell Metabolism, 24(4):608–615, 2016.

31. Li Yang, Wei-Hua Wang, Wei-Lin Qiu, Zhen Guo, Erfei Bi, and Cheng-Ran Xu. A single-cell transcriptomic analysis reveals precise pathways and regulatory mechanisms underlying hepatoblast differentiation. Hepatology, 66(5):1387–1401, 2017.

